# Hydrogel Capsule-Based Digital Quantitative Polymerase Chain Reaction

**DOI:** 10.1101/2023.03.29.534823

**Authors:** Zheng Lin Tan, Masato Yasuura, Yukichi Horiguchi, Hiroki Ashiba, Takashi Fukuda

**Author notes:** Corresponding to Takashi Fukuda and Zheng Lin Tan.

## Abstract

Droplet digital PCR (ddPCR) is accurate in nucleic acid quantification owing to its linearity and high sensitivity. Amplification of nucleic acid in droplets, however, is limited by the stability of droplets against thermal cycling. While the use of fluorinated oil or supplementation of surfactant could improve the stability of droplets, this process has also greatly increased the cost of ddPCR and limited post-PCR analysis. Here, we report a novel method known as gel capsule-based digital PCR (gc-dPCR) which includes method to prepare hydrogel capsules encapsulating PCR reaction mix, conducting PCR reaction, and readout by either qPCR system or fluorescence microplate reader. We have compared our method to existing methods, i.e., quantitative PCR or vortex ddPCR. Our approach results in higher fluorescence intensity compared to ddPCR suggesting higher sensitivity of the system. As hydrogel capsules are more stable than droplets in fluorinated oil throughout thermal cycling, all partitions can be quantified, thus preventing loss of information from low-concentration samples. Our approach should extend to all droplet-based PCR methods. Generally, our approach has greatly improved ddPCR by increasing droplets stability and sensitivity, and reducing the cost of ddPCR, which help to remove the barrier of ddPCR in settings with limited resources.

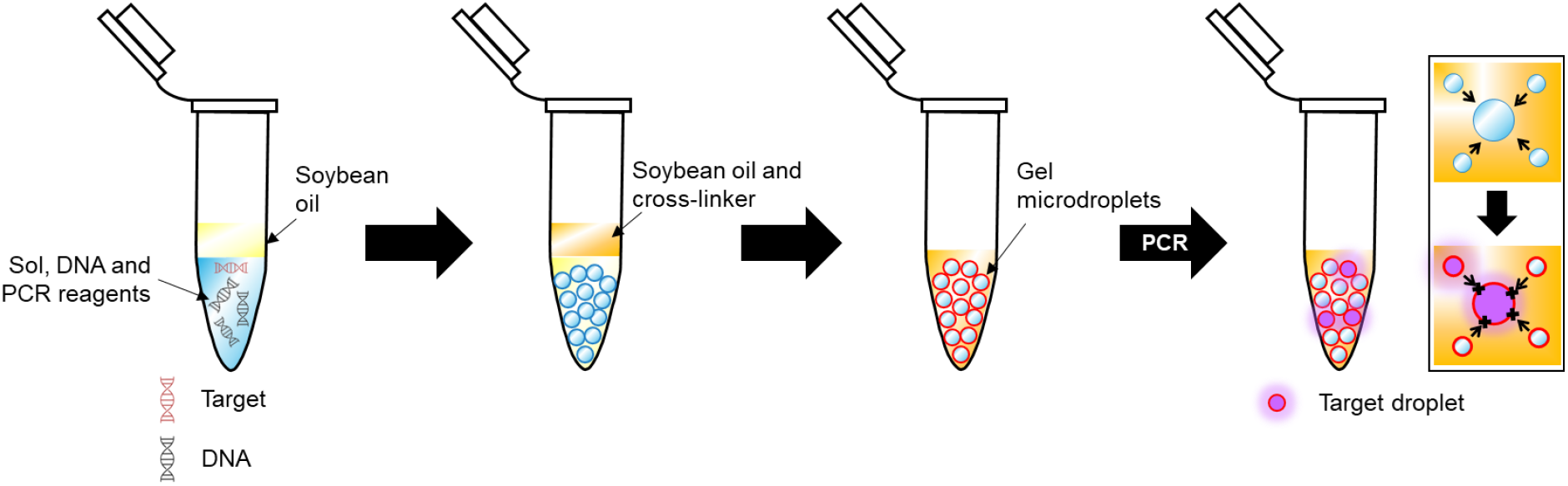

---

Technologies for nucleic acid quantification are important for science, industry, public health, and medical diagnosis. Quantitative polymerase chain reaction (qPCR) has emerged as the gold standard for its wide dynamic range and high specificity depends on probe[1, 2]. However, cumbersome optimization process is required for qPCR to obtain reliable and reproducible result[3]. In contrast, digital PCR (dPCR) based on limiting dilution of sample, such that each part of sample contains only one or no target molecule, detect sample after end-point amplification, enumerate number of positive parts of sample and yield concentration of target by statistics[4–6]. dPCR possess numerous advantages compared to qPCR, including absolute quantification without the needs of calibration curve, enhanced accuracy for small concentration (samples with quantification cycle, C_q_ ≥ 29)[7, 8], increased resistance against reaction inhibition[9], and ability to differentiate fragmented from intact molecules[10, 11], which make dPCR a valuable tool for clinical applications.

Droplet dPCR (ddPCR) is a variation of dPCR which rely on active emulsification method to compartmentalize samples into small volume of monodisperse droplets in oil[12, 13]. While ddPCR is exhibit an improved accuracy compared to other PCR approaches, requirement of equipment for active emulsification increases the cost of PCR and the barrier of adoption, which has limited the widespread of this technology. Furthermore, concerns remained for the stability of droplets throughout multiple thermal cycles required for nucleic acid amplification. While previous study has suggested that interaction of sodium bis(2-ethylhexyl) sulfosuccinate or polyglycerol polyricinoleate with bovine serum albumin at the interface of droplets could stabilize droplet during thermal cycle[14], use of proteins is not favorable, as autofluorescence from proteins might affect sensitivity of ddPCR system. To address this issue, fluorinated oil and fluorinated surfactant were used to reduce droplet coalescence during thermal cycling. Nevertheless, fluorinated surfactant does not completely prevent coalescence of droplets, i.e., fluorinated oils have densities between 1.4 – 1.9 g cm^-3^, e.g., FluoOil series by Emulseo has density ranging from 1.61 – 1.85 g cm^-3^ [15], which results in the floating of droplets on oil, and can cause aggregation and coalescence. Furthermore, global supply of fluorinated surfactant and fluorinated oil is limited, and the cost is high. For instance, 10 ml 2% fluorinated surfactant (FluoSurf by Emulseo) in fluorinated oil (Novec HFE 7500 by 3M) cost US$ 240.00 [16] (12^th^ January 2023). Furthermore, as the world leading manufacturer of fluorinated oil, has announced to exit production of fluorinated oil by 2025 due to environmental pollution[17], there is an urgent need to develop a novel method to replace the use of fluorinated oil for droplet stabilization. On the other hand, irreversible interfacial stability provided by fluorinated oil has limited the downstream application of droplets, e.g., next generation sequencing due to complexity in nucleic acid recovery. To further improve the stability of droplet, to reduce the cost of ddPCR, and to expand potential application of ddPCR by enabling post-ddPCR droplet manipulation, thus yield its benefit for wide range of applications, a new approach for fluorinated oil-independent droplets stabilization is required.

In this study, we demonstrate droplet stabilization with hydrogel. Rather than stabilization of phase interface, reversible solidification of reactor will provide higher mechanical and thermal stability to droplets. We mix the nucleic acid and PCR reagent with sodium alginate and crosslink sodium alginate gel with group 2 metal ion. We have selected barium as crosslinker against calcium. Whilst calcium has been widely used to crosslink alginate hydrogel, it is known to be a polymerase inhibitor. Compression of hydrogel capsules against 2 pieces of glass slides suggested that the hydrogel capsules is stable against mechanical forces. With this stability, we could generate and maintain droplet in soybean oil or mineral oil, which the price is < 10^−3^ – 10^−4^-fold of fluorinated oil with fluorinated surfactant, even after considering the price of alginate salt. We demonstrate this method by quantifying influenza A virus (IAV) nucleic acid in hydrogel capsules by vortex ddPCR (vddPCR)[18]. While we use vddPCR as an example, our method should apply to any dPCR approach in which the reactors can be sealed, polymerization can be easily reversed, and the samples can be easily recovered.

## Experimental

### Preparation of barium alginate hydrogel capsule

Barium chloride (Wako Pure Chemical, Cat # 023-00195, Osaka, Japan) was dissolved in soybean oil (Wako Pure Chemical, Cat # 190-03776, Osaka, Japan) at final concentration of 20 mmol L^-1^. While stirring, dropped 2% (w/v) sodium alginate (Wako Pure Chemical, Cat # 194-13321, Osaka, Japan) with or without red dye into soybean oil with barium chloride. The mixture was stirred for 60 s and observed with an optical microscope. To conduct qualitative analysis of compression of hydrogel capsule, hydrogel capsules were sandwiched between 2 pieces of glass slides. Then, stress was applied at the end of glass slide with a pair of clamps. The mechanical stability of hydrogel capsules was evaluated by observation of compression and leakage of content from hydrogel capsules with microscope when the diameter of hydrogel capsules increased by at least 10%.

### qPCR and Bulk Readouts of ddPCR

Hydrolysis probe based on fluorescein and carboxytetramethylrhodamine was used in qPCR. The TaqMan Assays were rigorously designed, and the specificity of these assays were determined by the vendor (Thermofisher Scientific). The reaction mixtures were prepared by mixing 4× TaqMan Fast Virus 1-Setp Master Mix (Thermofisher Scientific, Ref # 4444427, Vilnus, Lithuania), 10^7^ copies mL^-1^ influenza A virus standard DNA, primers and probes for matrix protein of IAV as in instruction. For qPCR and vddPCR assays, distilled deionized water (Nippon Gene, Cat # 318-90105, Toyama, Japan) was used as diluent, while 2% (w/v) sodium alginate solution in distilled deionized water (1% (w/v) final concentration) was used as diluent for gc-dPCR assays. For each assay, 20 μL reaction mix was used.

To generate droplets for ddPCR, reaction mix were vortex in 15 μL Automated Droplet Generation Oil for Probes (Bio-Rad, Cat # 1864110, USA) for 5 min, followed by addition of 15 μL Automated Droplet Generation Oil for Probes with barium chloride and vortex for another 5 min. For qPCR assays, the reaction mix were covered with 30 μL Automated Droplet Generation Oil for Probes with barium chloride to prevent measurement bias. These assays were incubated in qPCR system (Roche, LightCycler 96, Switzerland) as described in manufacturer’s instruction of 4× TaqMan Fast Virus 1-Setp Master Mix, and the C_q_ were recorded.

After incubation, the assays were transferred into a black, flat-bottom 96-well plate, and the fluorescence intensity with excitation/emission wavelength at 488/530 nm was measured with fluorescence microplate reader (Thermo Scientific, VarioSkan Lux, Singapore). For each sample, 6 replicates were prepared with at least 2 different aliquots of reagents to ensure reproducibility.

### Comparison of gc-dPCR in Fluorinated Oil and Soybean Oil

Hydrogel capsules were prepared with either fluorinated oil or soybean oil as described in previous section. Then, the hydrogel capsules were incubated in thermal cycler. After incubation, the fluorescence intensity with excitation/emission wavelength at 488/530 nm was measured with fluorescence microplate reader. The fluorescence intensity of equal volume of fluorinated oil and soybean oil were also measured to evaluate autofluorescence. After measurement, the hydrogel capsule from each condition was observed with fluorescence microscope (Olympus, BX3-RFAS, China; objective lens: Olympus, UPFLN10X2, Tokyo, Japan; fluorescence mirror unit: Evident Corporation, U-FBW, Nagano, Japan; illuminator: Thorlabs, M455F1, New Jersey, USA; complementary metal-oxide semiconductor camera: Hamamatsu Photonics, C13440-20CU, Hamamatsu, Japan). For each sample, 11 replicates were prepared with at least 2 different aliquots of reagents. Two measurements were conducted on different day with 6 replicates on the one day and 5 replicates on another day to ensure reproducibility.

### Detection of Influenza A Virus RNA with gc-dPCR

To ensure that the proposed method is applicable in virus detection, nucleic acid was extracted from sample with unknown concentration of influenza A virus (A/Puerto Rico/8/1934(H1N1)). The nucleic acid extract was diluted by 3000-fold. Reaction mixtures were prepared as described in previous section, except influenza A virus standard DNA was replaced with nucleic acid extract. Mixture without sample and mixture without primers were used as control. Hydrogel capsules was generated with soybean oil as described in previous section. RNA was amplified as manufacturer’s instruction for 45 cycles. Both C_q_ and endpoint fluorescence intensity were measured. For each sample, 3 independent tests were conducted.

## Results and Discussion

ddPCR is more sensitive than qPCR due to its nature of linearity, which allow it to accurately quantify nucleic acid, particularly at low copy number. However, high temperature will decrease viscosity of oil and increase kinetic energy of droplets, which results in increased probability of droplet coalescence. Condensation of droplets increases surface charge, which attract small droplets, and further increased the probability of coalescence. While it has been demonstrated that fluorinated oil with fluorinated surfactant could increase stability of droplets throughout PCR, some properties, e.g., high density, low surface tension, and high evaporation rates of fluorinated oil[19] will expose droplets to air, and result in evaporation of droplets during PCR, in addition to its high cost. Furthermore, stability of fluorinated oil at low temperature limits demulsification of droplets, thus limited further analysis of post-PCR products. By adding sodium alginate into reaction mix, and polymerize droplets divalent ions[20], it is possible to stabilize droplets without fluorinated oil, while increasing mechanical stability of droplets (Figure 1). In our study, barium was used instead of other divalent ions as these ions are PCR inhibitors[21]. Furthermore, conventional calcium alginate is sensitive to the presence of phosphate, citrate, sodium, and magnesium which will result in gel swelling followed by leakage of content.

**Figure 1.**
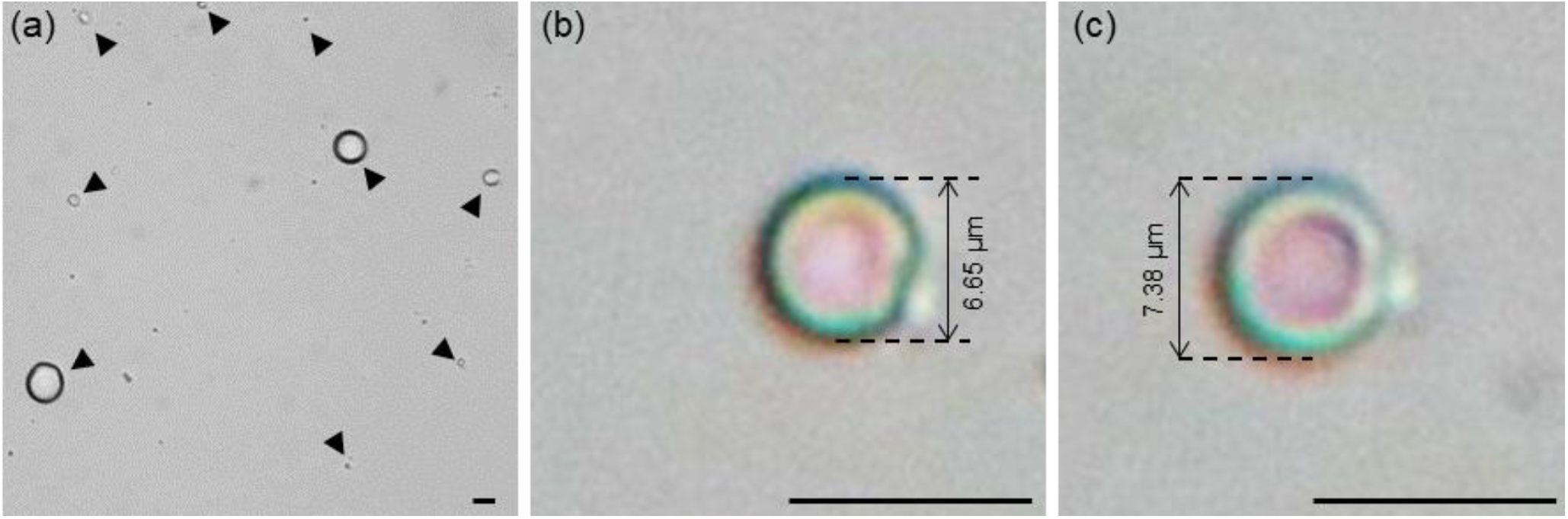
Mechanical stability of barium alginate hydrogel capsules. (a) barium alginate hydrogel capsules generated with soybean oil. Black arrows show the positions of gel microdroplets. Mechanical stability of gel against mechanical stress was evaluated by micrographs of hydrogel capsules encapsulating red dye (b) before and (c) during compression. Expansion of droplets due to compression was compared with black dashed line. The scale bars are 10 μm.

Furthermore, we have also demonstrated that barium alginate is stable against mechanical stress. Compression has resulted in increased in diameter of hydrogel capsules from 6.65 μm to 7.38 μm. Despite of the deformation, no leakage of dye from hydrogel capsule was observed (Figure 1b, c), which suggested that hydrogel capsule is stable against mechanical stress. Moreover, a layer of shell-like structure was observed under high magnification when the hydrogel capsule was compressed, which suggested that surface polymerization might have occurred, and formed a barium alginate gel shell bead. Mørch et al. has investigated the distribution of alginate after cross-linking with various types of type 2 metal ions. Their study suggested that with low concentration of type 2 metal ions[22], i.e., ≤20 mmol L^-1^, a rapid gelling could occur, which results in accumulation of alginate at the outer layer. In our study, we have provided evident that a gel shell might have formed at the outer layer of gel microdroplets based on the difference of color between core and outer layer, which stemmed from the difference in optical properties of gel and sol, and their response toward mechanical stress. Formation of gel shell bead with liquid core is more favorable than complete polymerization of alginate hydrogel as lower density of gel core could facilitate rapid reaction compared to solid core. Furthermore, supplementation of chelating agent, i.e., ethylenediaminetetraacetic acid to barium alginate gel could dissolved the gel shell (Figure S1) of gel microdroplets, which will enable post-analysis manipulation of droplets.

Recently, Abate’s group has demonstrated accurate quantitation of severe acute respiratory syndrome coronavirus 2 sequences with vddPCR[18]. The process is convenient as it does not required microfluidics, and it can be conducted entirely in bulk processing, which the size distribution of droplets is not an important parameter. Furthermore, the quantitative property of vddPCR was also demonstrated by Abate’s group. In this study, we have compared our proposed method, gel capsule-based dPCR (gc-dPCR) to vddPCR. To ensure that all reactions were conducted under similar conditions, fluorinated oil with barium chloride was supplemented to all assays, either to cover the reaction mix or as continuous phase to disperse droplets. Prior study has described that dispersion of reaction mix, even with vortex emulsification could reduce C_q_ of qPCR[18]. Our result has confirmed the effect of compartmentalization in reducing C_q_ of qPCR (Figure 2a). Endpoint measurement with fluorescence microplate reader has shown significant increase in fluorescence intensity in vddPCR and gc-dPCR assay compared to qPCR assay, while no statistical significant was observed between vddPCR and gc-dPCR assay (Figure 2b). Considered both the C_q_ and fluorescence intensity, although supplementation of hydrogel into reaction mix has significantly increased C_q_ of qPCR, however, it did not increase the fluorescence intensity, suggesting that the difference in C_q_ is not practical significant. These results suggested that both vddPCR and gc-dPCR could increase sensitivity of qPCR.

**Figure 2.**
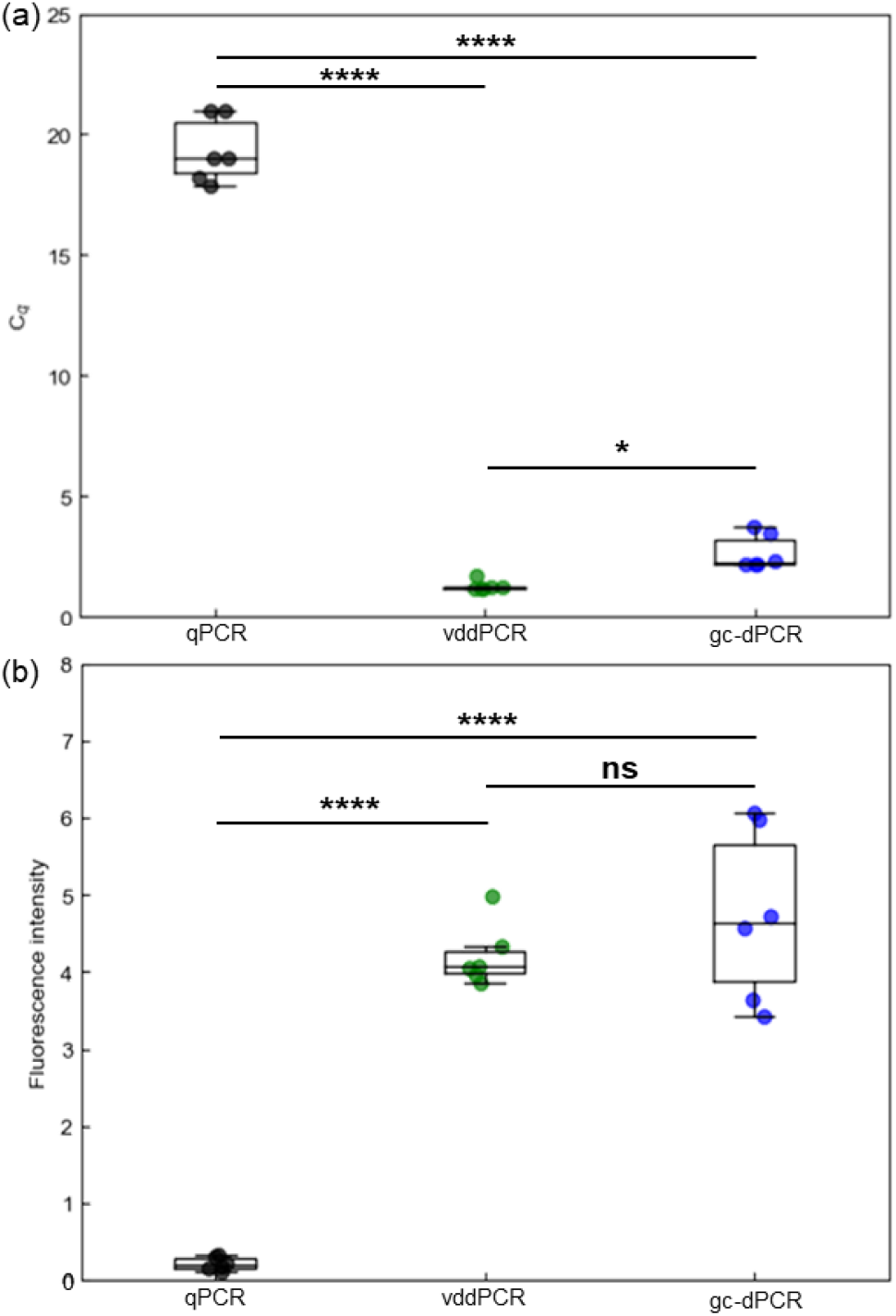
Gellification of droplets retain the efficiency of vddPCR. (a) qPCR readout of vddPCR and gc-dPCR suggested that both vddPCR and gc-dPCR are more sensitive than qPCR. (b) Endpoint measurement of each assay. vddPCR and gc-dPCR produced stronger fluorescence intensity compared to qPCR. The boxplots represent statistical analysis of data obtained from 6 independent tests. Statistical significance was evaluated by one-way analysis of variance followed by post-hoc Tukey honest significance difference test. * *p* < 0.05, ** *p* < 0.01, *** *p* < 0.001, **** *p* < 0.0001. ns indicates not significant.

While it is generally regarded that higher viscosity of hydrogel (> 80 mPa s) compared to water (10 mPa s) will result in retarded reaction rate, in our study, PCR efficiency remained the same for both vddPCR and gc-dPCR as diffusion length of reactant in droplets is short enough to overcome the effect of viscosity. On the other hand, larger variability of C_q_ and fluorescence intensity were observed for gc-dPCR, which might be due to larger size distribution of hydrogel capsules resulted from higher viscosity of sodium alginate sol compared to water. Nevertheless, higher viscosity of sodium alginate sol will result in generation of microdroplets with larger size, which will result in production of more PCR products per target molecule, thus increase the sensitivity of system. Higher sensitivity and signal intensity from microdroplets with larger size has compensated the viscosity of sodium alginate sol, which results in similar efficiency between vddPCR and gc-dPCR. Another advantage of gc-dPCR is its capability to maintain its stability without the needs of fluorinated oil or surfactant. However, it remained uncertain whether hydrogel capsules generated with soybean oil without surfactant can be used in ddPCR. To evaluate the effect of oil on gc-dPCR, endpoint measurement PCR products after thermal cycling of hydrogel capsules generated with fluorinated oil and soybean oil were conducted (Figure 3).

**Figure 3.**
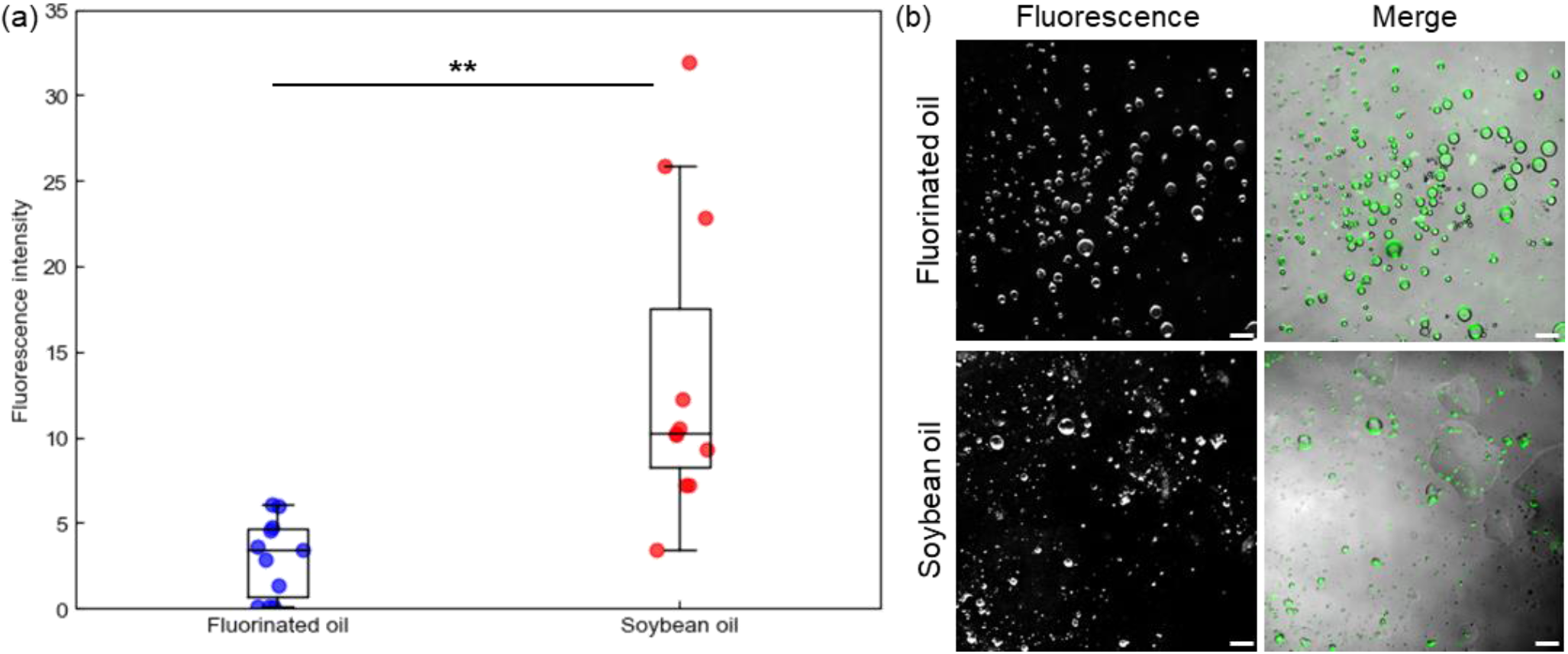
gc-dPCR in soybean oil. (a) The effect of oil on gc-dPCR. The boxplots represent statistical analysis of data obtained from 11 independent tests. Statistical significance was evaluated by Student’s t-test. * *p* < 0.05, ** *p* < 0.01. (b) Micrographs of hydrogel capsules after gc-dPCR. The hydrogel capsules from each condition was observed under fluorescence filter and merged image of brightfield and fluorescence micrographs. Green in merge channel was signal originated from fluorescein in fluorescence channel. The scale bars are 100 μm.

Although the fluorescence intensity of PCR products encapsulated in hydrogel capsules generated with soybean oil was higher than those encapsulated in hydrogel capsules generated with fluorinated oil, the variance was also larger (Figure 3a). The difference of variance of fluorescence intensity might be resulted from larger droplet size dispersity for droplets generated with soybean oil compared to fluorinated oil (Figure 3b). Previous studies have experimentally shown that viscosity of continuous phase (oil) is a parameter of droplet size and its distribution[23, 24]. Higher viscosity of continuous phase will result in larger size dispersity of droplets. Consider the viscosity of soybean oil at 25°C is 50.09 mPa s, which is 12.5 – 41.7-fold higher than fluorinated oil (1.2 – 4 mPa s), thus, hydrogel capsules generated with soybean oil will have larger dispersity compared to hydrogel capsules generated with fluorinated oil. The variance can be reduced by generating droplets with active emulsification methods which produce higher monodispersity of droplets.

On the other hand, the density of soybean oil (0.917 g cm^-3^) approximates the density of water and lower than fluorinated oil (1.4 – 1.6 g cm^-3^). While barium alginate hydrogel capsules float on fluorinated oil, they distributed evenly in soybean oil (Figure S2), which results in efficient amplification of nucleic acid due to even heating of hydrogel capsules. Efficient amplification results in higher signal intensity, hence, the fluorescence intensity of PCR products from hydrogel capsules generated with soybean oil is higher than those generated with fluorinated oil.

In addition, thermal stability of hydrogel capsules was also demonstrated. Fluorophore remained encapsulated in capsule after PCR thermal cycling (Figure 3b), suggesting that no breakage or merger of capsule which will resulted in leakage of fluorophore has occurred. As the outer layer of hydrogel capsule is gellified, and the onset decompose temperature of barium alginate is 150°C[25], barium alginate capsule is stable within the range of temperature for thermal cycling.

Furthermore, the autofluorescence of fluorinated oil and soybean oil was confirmed (Table 1). Autofluorescence of soybean oil originated from fatty acid chain is a concern in fluorescence measurement. Although the autofluorescence intensity of soybean oil is higher than fluorinated oil by 2-fold, the autofluorescence is negligible compared to the fluorescence intensity generated from PCR products (Figure 3a). As fluorescent tag binds to nucleic acid, and the phosphate backbone of nucleic acid which expose to the external environment is hydrophilic, leakage of fluorescent tag into oil is negligible. These results suggested that higher fluorescence intensity from hydrogel capsules generated with soybean oil was the result of even distribution of droplets in oil.

**Table 1.**
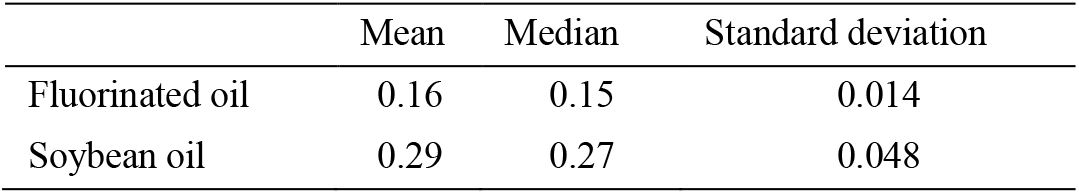
Autofluorescence of oils. Statistical analysis of fluorescence intensity of fluorinated oil and soybean oil from 3 independent tests.

To demonstrate the applicability of our method on actual sample, gc-dPCR was performed on actual sample containing unknown concentration of influenza A virus (Table 2). Increase in fluorescence intensity and decrease in C_q_ suggested that influenza A virus was detected, and our method is applicable in actual sample.

**Table 2.** Detection of RNA of influenza A virus from actual sample. Each set of data is obtained from 3 independent tests.

| Sample | C<sub>q</sub> |  |  | Fluorescence intensity |  |  |
| --- | --- | --- | --- | --- | --- | --- |
|  | Mean | Median | S.D. | Mean | Median | S.D. |
| Control without target RNA | N.D. | N.D. | N.D. | 0.22 | 0.16 | 0.13 |
| Control without primers | N.D. | N.D. | N.D. | 0.10 | 0.10 | 0.00 |
| Sample containing influenza A virus | 19.10 | 19.05 | 0.18 | 28.64 | 30.92 | 5.90 |

## Conclusions

Our approach is a major advance because it has greatly enhanced the stability of droplets, reduced the running cost of ddPCR by approximately 4000-fold through replacement of fluorinated oil with soybean oil and sodium alginate, and provided the option for post-PCR droplet manipulation. This method is compatible with various droplet generation approaches, e.g., microfluidic-based droplets generation, membrane emulsification and droplet generation in turbulent flow. While hydrogel increases the stability of microdroplets, viscosity of sol could also increase the pressure on membrane when membrane with small pore size was used in emulsification. For the case when smaller capsule size and monodisperse gel capsule is required to absolute quantification by positive capsule quantification, microfluidic device[26, 27] or low viscosity sol can be used. The stability provided by hydrogel capsules afford other valuable benefits, including the ability to alter the composition of continuous phase to ensure even distribution of droplets and heat conduction, reduce scattering at the edge of droplets, and post-quantification analysis of microdroplets of interest, e.g., isolating and sequencing of nucleic acid encapsulated in droplets. Furthermore, owing to the benefit of high mechanical stability of gel microdroplets, high-throughput fluorescence-activated cell sorter or fluorescence-activated droplet sorter can be used to analyze and isolate droplets[26, 27] of interest at throughput of > 500 kHz[28]. These features are particularly useful in clinical diagnosis and to investigate mutation in sequence. Moreover, even distribution of droplets in oil could ensure even heat transfer to all droplets, which results in even reaction. Even amplification reaction in turn increases the yield of amplicon (fluorescence intensity), which can help to reduce false positive test results by setting a higher threshold value. Hydrogel capsules have demonstrated high mechanical and thermal resistance without surfactant, which could greatly reduce the cost of ddPCR, and helps to accelerate employment of ddPCR method even in settings with limited resources.

## Acknowledgement

Not applicable

## Author Contributions

Conceptualization: TF and ZLT; Experimental design and data curation: ZLT, TF, MY; Data analysis: ZLT; Interpretation of results: ZLT, MY, TF, YH, HA; Funding and administration: TF, MY; Writing – Original draft: ZLT, MY, TF; Writing – Review and editing: HA, YH, TF, MY, ZLT

## Competing Interests

The authors declare no competing interests.

## Notes

### Competing Interest Statement

The authors have declared no competing interest.

### Summary of Updates

The term "hydrogel microdroplets" was replaced with "gel capsules"; "gel ddPCR" was replaced with "gel capsule-based dPCR" to avoid ambiguity. Result for additional experiment which involve application of our method in actual sample has also been included.

