## Supplementary material for "Hydrogel Capsule-Based Digital Quantitative Polymerase Chain Reaction": Figure S1; Figure S2

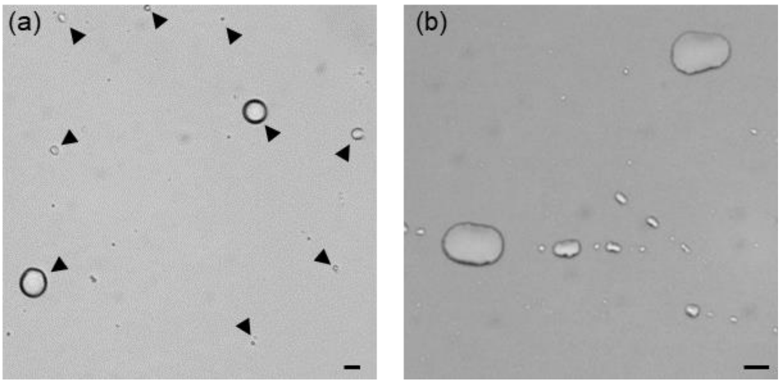


**Figure S1** Cross-linking of sodium alginate by barium chloride is reversible. Barium alginate gel microdroplets (a) before and (b) after ethylenediaminetetraacetic acid treatment. Ethylenediaminetetraacetic acid dissolved the gel shell of barium chloride, which results in further compression and coalescence of droplets. The scale bars are 10 µm.

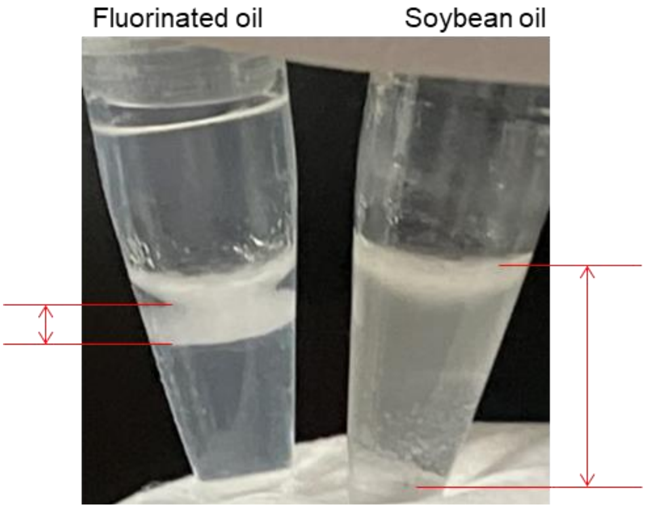


**Figure S2** Suspension of hydrogel microdroplets in fluorinated oil and soybean oil after thermal cycling. Region marks in red indicates the region where emulsion was suspended.
